# Chromosome-level genome assembly of the common chiton, *Liolophura japonica* (Lischke, 1873)

**DOI:** 10.1101/2024.01.15.575488

**Authors:** Hong Kong Biodiversity Genomics Consortium, Project Coordinator and Co-Principal Investigators, Jerome H.L. Hui, Ting Fung Chan, Leo L. Chan, Siu Gin Cheung, Chi Chiu Cheang, James K.H. Fang, Juan D. Gaitan-Espitia, Stanley C.K. Lau, Yik Hei Sung, Chris K.C. Wong, Kevin Y.L. Yip, Yingying Wei, DNA extraction, library preparation and sequencing, Franco M.F. Au, Wai Lok So, Genome assembly and gene model prediction, Wenyan Nong, Gene family annotation, Ming Fung Franco Au, Samples collectors, Tin Yan Hui, Brian K.H. Leung, Gray A. Williams

## Abstract

Chitons (Polyplacophora) are marine molluscs that can be found worldwide from cold waters to the tropics, and play important ecological roles in the environment. Nevertheless, there remains only two chiton genomes sequenced to date. The chiton *Liolophura japonica* (Lischke, 1873) is one of the most abundant polyplacophorans found throughout East Asia. Our PacBio HiFi reads and Omni-C sequencing data resulted in a high-quality near chromosome-level genome assembly of ∼609 Mb with a scaffold N50 length of 37.34 Mb (96.1% BUSCO). A total of 28,233 genes were predicted, including 28,010 protein-coding genes. The repeat content (27.89%) was similar to the other Chitonidae species and approximately three times lower than in the genome of the Hanleyidae chiton. The genomic resources provided in this work will help to expand our understanding of the evolution of molluscs and the ecological adaptation of chitons.

## Introduction

The Mollusca is the second largest animal phylum after Arthropoda, which divides into two subphyla, including the shell-bearing Conchifera (Monoplacophora, Bivalvia, Gastropoda, Scaphopoda and Cephalopoda) and the shell-lacking Aculifera (Polyplacophora, Caudofoveata and Solenogastres) (Kocot et al 2020; Kocot et al 2011; Smith et al 2011). Within the latter, chitons (Polyplacophora) are thought to be a relatively early diverging group of living molluscs (Sigwart and Sutton, 2007), represented by keystone species that regulate the structure of marine communities in both intertidal and subtidal systems worldwide (Paine, 2002; Martinez-Tejada et al., 2016). These “living fossils” are characterised by a highly evolutionary-conserved and unique type of shell formed by eight articulating aragonite plates that provide protection from environmental threats (Eernisse & Reynolds 1994; Scherholz et al 2013; Connors et al., 2012). This biomineralized armour incorporates an integrated sensory system that includes hundreds of eyes with aragonite- based lenses (Li et al., 2015) acting as a light-sensing adaptation (Chappell et al., 2023). Contrary to other molluscs, our understanding of the Polyplacophora is constrained to only two available genomes (*Acanthopleura granulata* and *Hanleya hanleyi*) (Varney et al 2021;2022

*Liolophura japonica* (Polyplacophora, Chitonidae) (Lischke 1873) (Figure 1A) is one of the most abundant polyplacophorans found on intertidal rocky reefs of the Asian continent including China, Korea, and Japan. This species diverged from the last common ancestor of *Liolophura* at ∼184 million years ago during the early Pleistocene period (Choi et al 2021). On the shore, they are distributed over a wide vertical range from the mid littoral zone to the low subtidal zone, where the animals experience periodic fluctuations in environmental conditions with the ebb and flow of the tide (Harper 1996; Harper & Williams 2001). Unlike other mobile species inhabiting rocky shores which migrate towards the low shore areas during summer (Lewis 1954; Williams & Morritt 1995), *L. japonica* does not show any significant seasonal migration behaviour and can survive stressful low tide periods presumably by fitting themselves into small refuges via the eight flexible, interlocking plates (Ng & Williams 2006; McMahon 1986; Harper & Williams 2001). In terms of its feeding biology, it is a generalist consumer feeding on a wide range of microalgae and macroalgae (Nishihama et al 1986; Harper 1996), and owing to their high density it is an important grazer in the intertidal zone of East Asia, contributing to the control of on-shore primary productivity (Dethier & Duggins 1984; Nishihama et al 1986).

**Figure 1.**
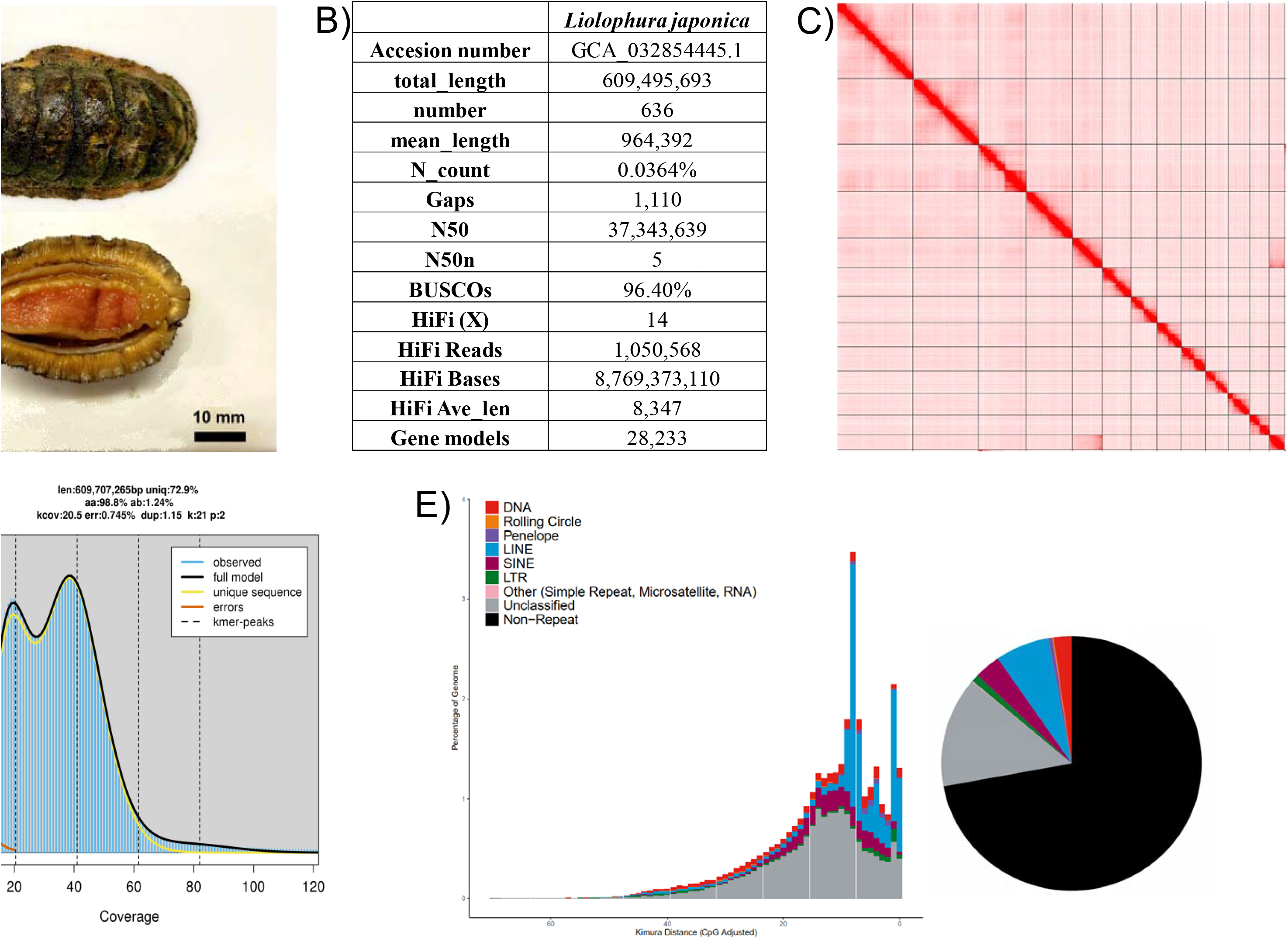
A) Picture of *Liolophura japonica*; B) Statistics of the genome assembly generated in this study; C) Hi-C contact map of the assembly visualised using Juicebox v1.11.08 (details can be found in Supplementary Table 6); D) Genomescope report with k-mer = 21; E) Repetitive elements distribution.

## Context

Here we report the assembled genome of the chiton *L. japonica* (Polyplacophora, Chitonidae) (Lischke 1873) (Figure 1A), which is selected as one of the species sequenced by the Hong Kong Biodiversity Genomics Consortium (a.k.a. EarthBioGenome Project Hong Kong), organised by researchers from 8 publicly funded universities in Hong Kong. The *L. japonica* genome presented in this study is of high quality and near chromosomal level, which serves as a valuable resource for the understanding of evolutionary biology of polyplacophorans as well as the adaptation of its resilience under oscillating environmental changes in the intertidal zones.

## Methods

### Collection and storage of samples, Isolation of high molecular weight genomic DNA, quantification, and qualification

The chiton, *Liolophura japonica*, were collected at the rocky shore in Kau Sai Chau, Hong Kong (22.380 °N, 114.310 °E) during the summer of 2022. They were kept at 35 ppt artificial sea water under room temperature until DNA isolation. High molecular weight (HMW) genomic DNA was isolated from single individual. In general, the tissue was first frozen in liquid nitrogen and ground to powder. DNA extraction was carried out with the sample powder using a Qiagen MagAttract HMW kit (Qiagen Cat. No. 67563) following the manufacturer’s protocol with some modifications. In brief, around 1 g of sample was first put in 200 μl 1X PBS and mixed with RNase A, Proteinase K, and Buffer AL provided in the kit. The mixture was allowed to sit at room temperature (∼22 °C) for 2 hours. The mixture was gently flicked to allow through mixing of samples and digestion solution. The sample was eluted with 120 μl of elution buffer (PacBio Ref. No. 101-633-500). When DNA was transferred at any time throughout the extraction progress, wide-bore tips were used. Afterwards, the sample was quantified by the Qubit^®^ Fluorometer and Qubit^™^ dsDNA HS and BR Assay Kits (Invitrogen^™^ Cat. No. Q32851). Overnight pulse-field gel electrophoresis was used to examine the molecular weight of the isolated DNA, together with three DNA markers (λ-Hind III digest; Takara Cat. No. 3403, DL15,000 DNA Marker; Takara Cat. No. 3582A and CHEF DNA Size Standard-8-48 kb Ladder; Cat. No. 170-3707). Purity of sample was examined by the NanoDrop^™^ One/OneC Microvolume UV-Vis Spectrophotometer, with A260/A280: ∼1.8 and A260/A230: >2.0 as a standard.

### DNA shearing, library preparation and sequencing

120 μl of DNA sample with around 8-10 μg DNA was transferred to a g-tube (Covaris Part No. 520079). The sample was then proceeded to 6 passes of centrifugation with 2,000 xg of 2 min. The resultant DNA was collected and stored in a 2 mL DNA LoBind^®^ Tube (Eppendorf Cat. No. 022431048) at 4 °C until library preparation. Overnight pulse-field gel electrophoresis was used to examine the molecular weight of the isolated DNA as described in the previous section. The electrophoresis profile was set as follow: 5K as the lower end and 100K as the higher end for the designated molecular weight; Gradient=6.0V/cm; Run time = 15 h:16 min; included angle = 120 °; Int. Sw. Tm = 22 s; Fin. Sw. Tm = 0.53 s; Ramping factor: a = Linear. The gel was run in 1.0% PFC agarose in 0.5X TBE buffer at 14 °C. A SMRTbell library was constructed using the SMRTbell® prep kit 3.0 (PacBio Ref. No. 102-141-700), following the manufacturer’s protocol. In brief, the genomic DNA was subjected to DNA repair to remove single-stranded overhangs and to repair damage on the DNA backbone during shearing. After repair, both ends of the DNA were polished and tailed with an A-overhang. Ligation of T-overhang SMRTbell adapters was performed at 20 °C for 30 min. Afterwards, the SMRTbell library was purified with SMRTbell^®^ cleanup beads (PacBio Ref. No. 102158-300). The concentration and size of the library were examined using the Qubit^®^ Fluorometer and Qubit™ dsDNA HS and BR Assay Kits (Invitrogen^™^ Cat. No. Q32851), and the pulse-field gel electrophoresis, respectively. A subsequent nuclease treatment step was performed to remove any non-SMRTbell structure in the library mixture. A final size-selection step was carried out to remove the small DNA fragments in the library with 35% AMPure PB beads. The Sequel^®^ II binding kit 3.2 (PacBio Ref. No. 102-194-100) was used for final preparation for sequencing. In brief, Sequel II primer 3.2 and Sequel II DNA polymerase 2.2 were annealed and bound to the SMRTbell library, respectively. The library was loaded at an on-plate concentration of 50-90 pM using the diffusion loading mode. The sequencing was conducted on the Sequel IIe System with an internal control provided in the binding kit. The sequencing was setup and performed in 30-hour movies (with 120 min pre-extension) with the software SMRT Link v11.0 (PacBio). HiFi reads are generated and collected for further analysis. One SMRT cell was used for this sequencing. Details sequencing data can be found in Supplementary Information 1.

### Omni-C library preparation and sequencing

An Omni-C library was constructed using the Dovetail® Omni-C® Library Preparation Kit (Dovetail Cat. No. 21005) according to the manufacturer’s instructions. In brief, around 20 mg of flash-freezing powered tissue sample was added into 1 mL 1X PBS, where the genomic DNA was crosslinked with formaldehyde and the fixed DNA was digested with endonuclease DNase I. Afterwards, the concentration and fragment size of the digested sample was checked by the Qubit® Fluorometer and Qubit™ dsDNA HS and BR Assay Kits (Invitrogen™ Cat. No. Q32851), and the TapeStation D5000 HS ScreenTape, respectively. Following the quality examination, both ends of the DNA were polished. Ligation of a biotinylated bridge adaptor was conducted at 22 °C for 30 min, and subsequent proximity ligation between crosslinked DNA fragments was performed at 22 °C for 1 hour. After the ligation events, the DNA was reverse crosslinked, and purified with SPRIselect™ Beads (Beckman Coulter Product No. B23317) to remove the biotin that was not internal to the ligated fragments. The Dovetail™ Library Module for Illumina (Dovetail Cat. No. 21004) was used for the end repair and adapter ligation. During this process, the DNA was tailed with an A-overhang, which allowed Illumina-compatible adapters to ligate to the DNA fragments at 20 °C for 15 min. The Omni-C library was then sheared into small fragments with USER Enzyme Mix and purified with SPRIselect™ Beads subsequently. Afterwards, Streptavidin Beads were added to isolate the DNA fragments with internal biotin. Universal and Index PCR Primers from the Dovetail™ Primer Set for Illumina (Dovetail Cat. No. 25005) were used to amplify the library. The final size selection step was carried out with SPRIselect™ Beads to pick only the DNA fragments ranging between 350 bp and 1000 bp. At last, the concentration and fragment size of the sequencing library was examined by the Qubit® Fluorometer and Qubit™ dsDNA HS and BR Assay Kits, and the TapeStation D5000 HS ScreenTape, respectively. After the quality checking, the library was sequenced on an Illumina HiSeq-PE150 platform. Details sequencing data can be found in Supplementary Information 1.

### Transcriptome sequencing

Total RNA and small RNA (<200 nt) from different tissues (i.e. digestive gland, foot, gill, gonad, heart) were isolated using the TRIzol reagent (Invitrogen) and the mirVana™ miRNA Isolation Kit (Ambion) following the manufacturer’s protocol respectively. The quality of the extracted RNA was checked using NanoDrop™ One/OneC Microvolume UV-Vis Spectrophotometer (Thermo Scientific™ Cat. No. ND-ONE-W) and gel electrophoresis. The qualified transcriptome samples were sent to Novogene Co. Ltd (Hong Kong, China) for polyA selected RNA sequencing library construction using the TruSeq RNA Sample Prep Kit v2 (Illumina Cat. No. RS-122-2001), and 150 bp paired-end (PE) sequencing. Agilent 2100 Bioanalyser (Agilent DNA 1000 Reagents) was used to measure the insert size and concentration of the final libraries. Details of the sequencing data are listed in Supplementary Information 1.

### Genome assembly, gene model prediction and repeat analysis

*De novo* genome assembly was performed using Hifiasm (Cheng et al 2021). Haplotypic duplications were identified and removed using purge_dups based on the depth of HiFi reads (Guan et al 2020). Proximity ligation data from the Omni-C library were used to scaffold genome assembly by YaHS (Zhou et al., 2022). Transposable elements (TEs) were annotated as previously described (Baril et al 2022), using the automated Earl Grey TE annotation pipeline (version 1.2, https://github.com/TobyBaril/EarlGrey). Mitochondrial genome was assembled using MitoHiFi (v2.2, https://github.com/marcelauliano/MitoHiFi) (Allio et al 2020). RNA sequencing data were first processed with Trimmomatic (Bolger, Lohse & Usadel 2014). Gene models were then trained and predicted by funannotate using the parameters “--repeats2evm --protein_evidence uniprot_sprot.fasta --genemark_mode ET --optimize_augustus --organism other --max_intronlen 350000”. Briefly, in the funannotate- train step, the transcripts assembled by Trinity were used to map to the repeat soft-masked genome by minimap2, the Trinity transcript alignments were converted to GFF3 format and used as input to run the PASA alignment in the Launch_PASA_pipeline.pl process to get the PASA models trained by TransDecoder, and then use Kallisto TPM data to select the PASA gene models. The PASA gene models were used to train Augustus in the funannotate-predict step. The gene models from several prediction sources including GeneMark, high-quality Augustus predictions (HiQ), pasa, Augustus, GlimmerHM and snap were passed to Evidence Modeler to generate the gene model annotation files. UTRs were captured in the funannotate-update step using PASA. Briefly, PASA was run twice in funannotate-update step, the transcripts generated in the previous funannotate-train step were mapped to the genome, these data were automatically parsed and used to update the UTR data using PASA comparison method according to PASA built-in process, 10,327 UTRs were updated in this study. The protein-coding genes were searched with BLASTP against the nr and swissprot databases by diamond (v0.9.24) (Buchfink, Xie & Huson, 2015) with parameters “--more-sensitive -- evalue 1e-3” and mapped by HISAT2 (version 2.1.0) (Kim et al., 2019) with transcriptome reads. 73.9% and 55.8% of the 28,032 protein-coding genes were mapped to the NR and swissprot databases, respectively. The BUSCO of the protein-coding genes was 90.9%, including 83.5% complete and single-copy genes, 7.4% complete and duplicated genes, 5.0% fragmented genes, and 4.1% of BUSCOs genes were missed (metazoa_odb10 with 954 total BUSCOs genes).

Transposable elements (TEs) were annotated as previously described (Baril et al 2022) using the automated Earl Grey TE annotation pipeline (version 1.2, https://github.com/TobyBaril/EarlGrey) with “-r eukarya” to search the initial mask of known elements and other default parameters. Briefly, this pipeline first identified known TEs from Dfam withRBRM (release 3.2) and RepBase (v20181026). De novo TEs were then identified and consensus boundaries extended using an automated BLAST, Extract, Extend process with 5 iterations and 1000 flanking bases added in each round. Redundant sequences were removed from the consensus library before the genome assembly was annotated with the combined known and de novo TE libraries. Annotations were processed to remove overlap and defragment annotations prior to final TE quantification.

## Results and discussion

A total of 8.77 Gb of HiFi bases of common chiton *Liolophura japonica* were generated with an average HiFi read length of 8,347 bp with 14X data coverage. After scaffolding with ∼397 Gb Omni-C data, the assembled genome size was 609.5 Mb, with 632 scaffolds and a scaffold N50 of 37.34 Mb, and the complete BUSCO estimation to be 96.1 % (metazoa_odb10) (Figure 1B; Table 1). 13 pseudomolecules of chromosomal length were anchored from Omni-C data (Figure 1C; Table 2), which is close to the karyotype of *L. japonica* (2n = 24)(Kawai 1976; Nishikawa & Ishida 1969) and thus indicates the assembly is of near chromosome-level. The assembled *L. japonica* genome has a genome size close to the estimation performed by GenomeScope, which is 609.7 Mb with heterozygosity rate at 1.24% (Figure 1D; Supplementary Information 2), and similar to the other published chiton genome (*Acanthopleura granulata*, 606 Mb) (Varney et al 2021) (Table 1). Telomeres can also be found in 7 out of 13 pseudomolecules (Table 3).

Total RNA sequencing data from different tissues, including digestive gland, foot, gill, gonad, and heart, was used to assemble the transcriptome of *L. japonica*. The final transcriptome assembly contained 294,118,260 transcripts, with 192,010 Trinity annotated genes (average length of 1,100 bp and N50 length of 2,373 bp). The resultant transcriptomes were used to predict the gene models, and a total of 28,233 gene models were generated with 28,010 predicted protein-coding genes, having a mean coding sequence length of 447 bp (Table 1).

For the repeat elements, a total repeat content of 27.89% was found in the genome assembly including 14.04% unclassified elements, comparable to the estimated genome repeat length with kmer 21 (27.14%) and the Chitonidae species *Acanthopleura granulata* (23.56%) (Varney et al 2021) (Figure 1E; Table 4; Supplementary Information 2). Among the remaining repeats, LINE is the most abundant (6.70%), followed by SINE elements (3.24%) and DNA (2.21%), whereas LTR, Penelope, rolling circle and other are only present in low proportions (LTR: 0.95%, Penelope: 0.52%, rolling circle: 0.15%, other: 0.08%).

## Conclusion and future perspective

The high-quality, near chromosome-level *L. japonica* genome was presented in this study. It provides a useful resource for further insights into the environmental adaptations of *L. japonica* in residing the intertidal zones and for future investigations on the evolutionary biology in polyplacophorans and other molluscs.

## Data validation and quality control

To assess the quality of samples in DNA extraction and PacBio library preparation, NanoDrop^™^ One/OneC Microvolume UV-Vis Spectrophotometer, Qubit^®^ Fluorometer, and overnight pulse-field gel electrophoresis were performed. Furthermore, the Omni-C library quality was assessed by Qubit® Fluorometer and TapeStation D5000 HS ScreenTape.

During genome assembly, the Hifiasm output was compared to the NT database via BLAST, which was used as the input for BlobTools (v1.1.1) (Laetsch & Blaxter 2017), where scaffolds recognised as possible contaminations were removed from the assembly (Supplementary Information 3). Moreover, a kmer-based statistical approach was employed to estimate genome heterozygosity, repeat content and genome size from sequencing Omni-C reads using Jellyfish (Marçais & Kingsford 2011) and GenomeScope (http://qb.cshl.edu/genomescope/genomescope2.0/) (Ranallo-Benavidez et al 2020) (Figure 1D, Supplementary Information 2). In addition, telomeric repeats was screened by using FindTelomeres (https://github.com/JanaSperschneider/FindTelomeres). Benchmarking Universal Single-Copy Orthologs (BUSCO, v5.5.0) (Manni et al 2021) was used to assess the completeness of the genome assembly and gene annotation with metazoan dataset (metazoa_odb10).

## Data availability

The final assemblies, Omni-C data and PacBio HiFi reads were submitted to NCBI under accession numbers GCA_032854445.1 (https://www.ncbi.nlm.nih.gov/datasets/genome/GCA_032854445.1/). The raw reads generated in this study were deposited to the NCBI database under the BioProject accessions PRJNA973839 (https://www.ncbi.nlm.nih.gov/bioproject/PRJNA973839). The genome annotation files were deposited and publicly available in figshare (https://doi.org/10.6084/m9.figshare.24208719).

## Author’s contributions

JHLH, TFC, LLC, SGC, CCC, JKHF, JDG, SCKL, YHS, CKCW, KYLY and YW conceived and supervised the study; MFFA and WLS carried out DNA extraction, library preparation and sequencing; WN performed genome assembly and gene model prediction; MFFA conducted homeobox gene annotation. TYH, BKHL and GAW collected the chiton samples.

## Funding

This work was funded and supported by the Hong Kong Research Grant Council Collaborative Research Fund C4015-20EF, CUHK Strategic Seed Funding for Collaborative Research Scheme (3133356) and CUHK Group Research Scheme (3110154).

## Table and figure legends

**Table 1.** Summary of genome statistics of *Liolophura japonica, Acanthopleura granulate* (Varney et al 2021) and *Hanleya hanleyi* (Varney et al 2022).

**Table 2.** Scaffold information of 13 pseudomolecules.

**Table 3.** List of telomeric repeats found in 9 scaffolds.

**Table 4.** Catalogue of repeat elements in the *Liolophura japonica* genome.

**Supplementary Information 1**. Details of genome and transcriptome sequencing data.

**Supplementary Information 2**. GenomeScope statistics report at K-mer = 21.

**Supplementary Information 3**. Genome assembly QC and contaminant/cobiont detection for the *Liolophura japonica*.

## Notes

### Competing Interest Statement

The authors have declared no competing interest.

