## Supplementary material for "Chromosome-level genome assembly of the common chiton, *Liolophura japonica* (Lischke, 1873)": EBPHK_Chiton_Supplementary Information.docx

**Supplementary Information 1.** Details of genome and transcriptome sequencing data.

| **Genome sequencing data** | | | | |
| --- | --- | --- | --- | --- |
| **Liabrary** | **No. of reads** | **No. of bases** | **Accesion** | **Coverage(X)** |
| PacBio HiFi | 1,050,568 | 8,769,373,110 | SRX20411988 | 14 |
| Omnic | 264,637,334 | 39,695,600,100 | SRX21911526 | 65 |
| **Transcriptome sequencing data** | | | | |
| **Sample name** | **No. of reads** | **No. of bases** | **Accesion** | **Tissue type** |
| Lj2HS_Dg_T | 35,899,054 | 5,384,856,396 | SAMN35319765 | digestive gland |
| Lj2HS_Ft_T | 35,289,414 | 5,293,410,229 | SAMN35319766 | foot |
| Lj2HS_Gl_T | 34,240,482 | 5,136,070,054 | SAMN35319767 | gill |
| Lj2HS_Gn_T | 31,128,962 | 4,669,342,254 | SAMN35319768 | gonad |
| Lj2HS_Ht_T | 37,837,458 | 5,675,616,543 | SAMN35319769 | heart |

**Supplementary Information 2.** GenomeScope statistics report at K-mer = 21.

| **Property** | **min** | **max** |
| --- | --- | --- |
| Homozygous (aa) | 98.74% | 98.78% |
| Heterozygous (ab) | 1.22% | 1.26% |
| Genome Haploid Length (bp) | 608,187,534 | 609,707,265 |
| Genome Repeat Length (bp) | 165,056,812 | 165,469,254 |
| Genome Unique Length (bp) | 443,130,722 | 444,238,011 |
| Model Fit | 78.27% | 99.09% |
| Read Error Rate | 0.74% | 0.74% |

**Supplementary Information 3.** Genome assembly QC and contaminant/cobiont detection for the *Liolophura japonica*.

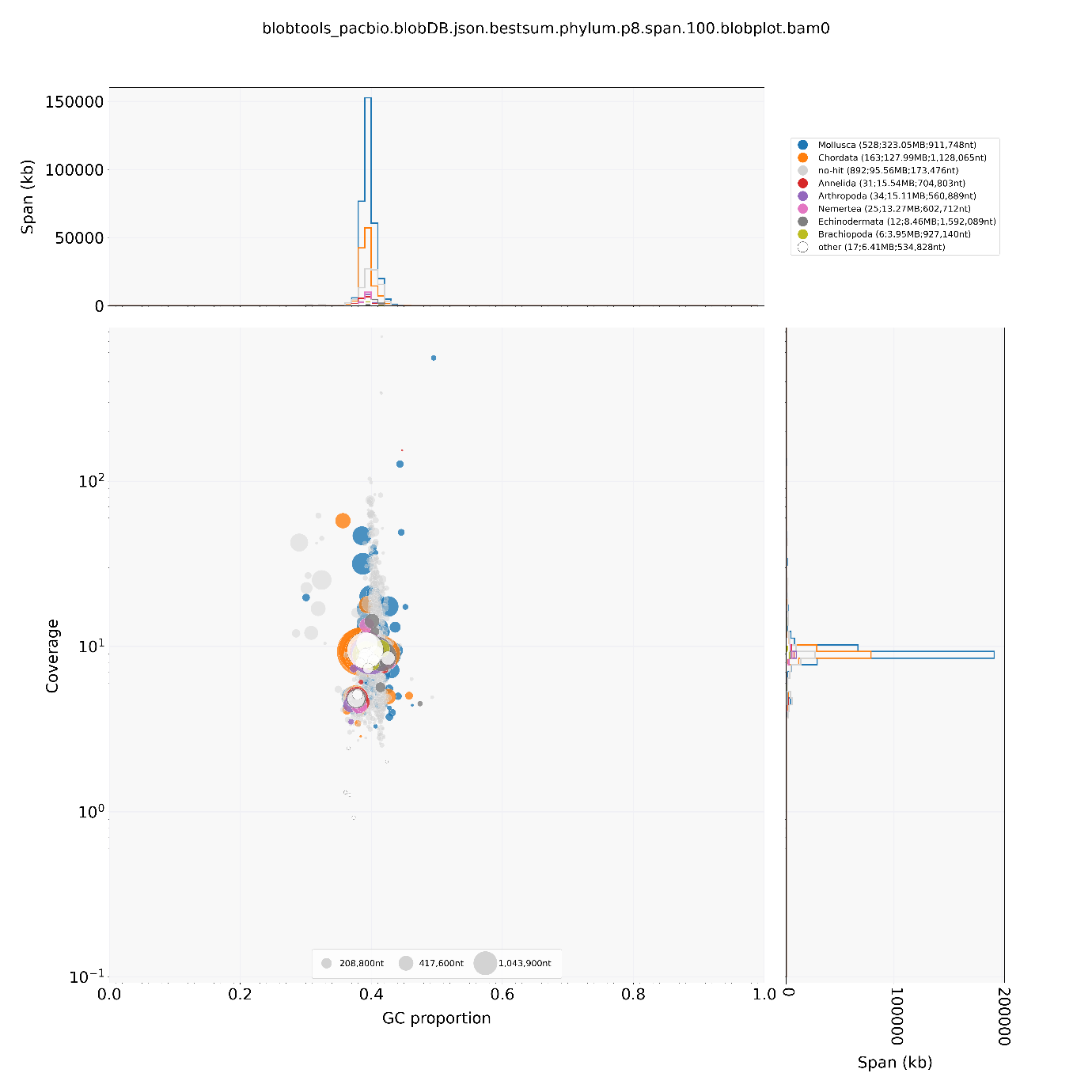

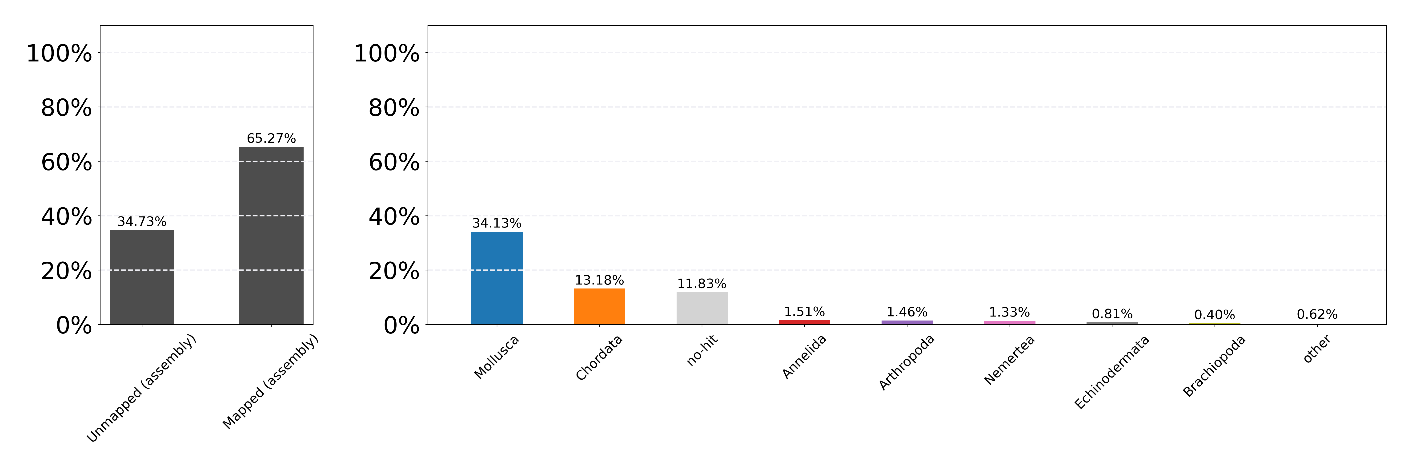
